# Anti-cancer properties of cannabidiol and Δ9-tetrahydrocannabinol and potential synergistic effects with gemcitabine, cisplatin and other cannabinoids in bladder cancer

**DOI:** 10.1101/2021.03.25.436633

**Authors:** Andrea M. Tomko, Erin G. Whynot, Denis J. Dupré

**Author notes:** **Correspondence:** Denis J. Dupré, 5850 College St., PO BOX 15 000, Sir Charles Tupper Medical Building, Faculty of Medicine, Dept. Pharmacology, Halifax, NS, B3H 4R2, Canada.

## Abstract

**Introduction:** With the legalization of cannabis in multiple jurisdictions throughout the world, a larger proportion of the population consumes cannabis. Several studies have demonstrated anti-tumor effects of components present in cannabis in different models. Unfortunately, little is known about the potential anti-tumoral effects of cannabinoids in bladder cancer, and how cannabinoids could potentially synergize with chemotherapeutic agents. Our study aims to identify whether a combination of cannabinoids, like cannabidiol and Δ9-tetrahydrocannabinol with agents commonly used to treat bladder cancer, such as gemcitabine and cisplatin, is able to produce desirable synergistic effects. We also evaluated whether co-treatment of different cannabinoids also generated synergistic effects.

**Materials and Methods:** We generated concentration curves with different drugs to identify the range at which they could exert anti-tumor effects. We also evaluated the activation of the apoptotic cascade and whether cannabinoids have the ability to reduce invasion.

**Results:** Cannabidiol, Δ9-tetrahydrocannabinol and other cannabinoids reduce cell viability of bladder cancer cell lines, and their combination with gemcitabine or cisplatin may induce differential responses: from antagonistic to additive and synergistic effects, depending on the concentrations used. Cannabidiol and Δ9-tetrahydrocannabinol were also shown to induce caspase 3 cleavage and reduce invasion in a Matrigel assay. Cannabidiol and Δ9-tetrahydrocannabinol also display synergistic properties with other cannabinoids like cannabichromene or cannabivarin.

**Discussion:** Our results indicate that cannabinoids can reduce human bladder transitional cell carcinoma cell viability, and that they can potentially exert synergistic effects when combined with other agents. Our *in vitro* results will form the basis for future studies *in vivo* and in clinical trials for the development of new therapies that could be beneficial for the treatment of bladder cancer in the future.

## Introduction

The most frequently encountered bladder cancer is transitional cell carcinoma (TCC), accounting for more than 90% of all bladder cancers (1). Lower grade, superficial non-muscle invasive tumours account for the majority of newly diagnosed TCC but a majority of tumours will recur in patients with worsening grade and stage (2). Without treatment, the median survival time before the development of effective chemotherapy rarely exceeded 3 to 6 months but advances in combination chemotherapy has increased median survival times to 14 months. Systemic combination chemotherapy, such as the methotrexate, vinblastine, doxorubicin and cisplatin (MVAC) regimen, has proven activity in advanced bladder cancer, but significant toxicity is observed with a treatment-related mortality of about 4%, and some patients are not eligible to receive cisplatin chemotherapy (3,4). Thus, there is a need for alternative therapies that provide improved survival outcomes or similar survival benefits with reduced toxicity compared to the MVAC regimen. Gemcitabine-based therapy can be used as intravesical instillations, with minimal bladder irritation, or systemic administration (5,6). Gemcitabine-cisplatin combination therapy is effective and safe and is frequently used as first-line therapy against metastatic bladder cancer (7–9). While the toxicity profile has been improved using this combination, the efficacy of the treatment remains relatively similar to treatment with the MVAC regimen.

Tobacco smoking is one of the most important risk factors for the development of bladder cancer and is associated with a 2-to 6-fold increase in the lifetime risk of urothelial cancer (10,11). Several studies have noted that a significant proportion of tobacco smokers also use cannabis. A study on the effects of cannabis and/or tobacco use was performed where men were followed over an 11-year period. Consumption of tobacco only was associated with an increased risk of bladder cancer (hazard regression [HR], 1.52), whereas cannabis use alone was associated with a 45% reduction in bladder cancer incidence (HR, 0.55). Using both cannabis and tobacco was associated with an intermediate HR of 1.28 (12). The metabolism of cannabis reveals that 65% of cannabis is excreted in the feces and 20% in urine (13). Chronic cannabis use causes accumulation of Δ9-tetrahydrocannabinol (THC) and its metabolite THC-COOH in adipose tissue such that it is excreted into the urine for as long as 30 to 60 days from the time chronic use is halted and it can be found in urine at levels greater than 500 ng/mL for chronic and/or recent Cannabis users.

Over 100 phytocannabinoids have been identified (14), but Δ9-tetrahydrocannabinol is the most common cannabinoid produced in the *Cannabis* plant (15). Cannabidiol is the most common cannabinoid in hemp and second most prevalent in the majority of cannabis cultivars, with a versatile pharmacological profile (16). Interestingly, studies have found that cannabinoids inhibit tumor cell growth and induce apoptosis in various cancer cells (17–23). Despite the use of cannabis in the population and evidence of antitumoral activity by cannabinoids, little is known about the anti-cancer effects of cannabis use in bladder cancer. Recently, a study suggested that cannabis-derived cannabichromene and Δ9-tetrahydrocannabinol displayed some synergy when used together in a model of urothelial cell carcinoma (24). Further research is required to understand the effect of the numerous compounds present in Cannabis to understand which ones exert the best antitumoral effects and how they may affect current chemotherapeutic agents. Our study presents the results of our investigation of the effects of Δ9-tetrahydrocannabinol and cannabidiol alone or in presence of other cannabinoids, gemcitabine, cisplatin or the combination of cisplatin and gemcitabine together in bladder cancer cell lines.

## Materials and Methods

### Drugs

Gemcitabine, cisplatin, Δ9-tetrahydrocannabinol and cannabidiol were obtained from Millipore-Sigma. The other cannabinoids were obtained from Toronto Research Chemicals.

### Cell Culture

Human Bladder transitional cell carcinoma T24 (ATCC^®^ HTB4™) and TCCSUP (ATCC^®^ HTB-5™) cells were cultured in McCoy’s 5A and Eagle’s Minimum Essential Medium (Millipore-Sigma), respectively, with 1% Penicillin-Streptomycin containing 10% fetal bovine serum (Gibco, Life Technologies) at 37°C, in a 5% CO_2_ atmosphere. It was demonstrated that *in vitro* models can adequately reproduce clinically relevant results and may be suitable to identify novel substances for the treatment of bladder cancer (25).

### Cytotoxicity Assays

Cells were seeded at 3,000 cells/well in 96-well plates and grown for 24 h before adding drugs. Cells were treated with increasing concentrations of Δ9-tetrahydrocannabinol, cannabidiol, gemcitabine and/or cisplatin for 48 h. To assess viability, AlamarBlue^®^ (Bio-Rad Laboratories) was added to each well and incubated for 4 h at 37 °C as per the manufacturer’s instructions. Fluorescence was measured following excitation at 540 nm and emission was read at 590 nm with a Biotek Cytation 3. Data are expressed as the percentage of viable cells vs. vehicle treated cells, normalized as 100% and represented as mean ± SEM. The *p* values represent data from at least three independent experiments.

### Cell Lysis and Western Blotting

Cells were lysed with RIPA buffer (150 mM NaCl, 50 mM Tris–HCl pH 7.5, 1% NP4O, 0.5% sodium deoxycholate, 0.1% sodium dodecyl sulfate, and 1 complete EDTA-free protease inhibitor cocktail tablet (Roche). BSA-coated beads (Protein A-Sepharose, Sigma-Aldrich) and 10% DNase I (Sigma-Aldrich) were added to remove nucleic acid and organellar material from the sample. Lysates were mixed 50:50 with 2X Laemmli Buffer and 2-mercaptoethanol (Bio-Rad Laboratories). Samples were run on a SDS–PAGE gel and transferred to nitrocellulose membranes before being blocked in a 10% skim milk powder/PBS solution for 60 min, and incubated overnight at 4 °C with their respective primary antibodies (cleaved Caspase 3 (p11): sc-271759 from Santa Cruz Biotechnologies). Chemiluminescence was performed on nitrocellulose membranes using Western Lightning® Plus-ECL Enhanced Chemiluminescence Substrate (PerkinElmer) before exposing them to X-ray film and development.

### Apoptosis Assay

Cells were grown on glass coverslips in 6 well plates and then treated with methanol or 2.5 μM Δ9-tetrahydrocannabinol and cannabidiol for 24 h. The Annexin V apoptosis detection kit (Santa Cruz Biotechnologies) was used to determine the rate of apoptosis. Cells were harvested and washed with PBS, then resuspended in Annexin V Assay Buffer following the manufacturer’s instructions. Cells were gently shaken in the dark with propidium iodide (PI) and Annexin V-FITC-conjugated stain for 20 min. Cells were then examined by fluorescence microscopy and at least 5 fields of view were recorded using an Olympus IX81 microscope equipped with a Photometrics coolSNAP HQ2 camera and an Excite series 120Q light source. Annexin V stain was excited at 488 nm and images were captured at 525 nm. PI was excited at 535 nm and images captured at 617 nm. Rates of early apoptosis were determined by dividing the number of cells that stained positive for Annexin-V divided by the total number of cells (26,27).

### Transwell Migration Assays

T24 cell cultures were prepared such that 5.0 × 10^5^ cells/condition were resuspended in DMEM and plated into the top portion of a transwell migration plate that contains a polycarbonate membrane with a pore size of 5.0 μm (Costar). In the bottom portion of the well, 600 μL of DMEM containing 10 ng/mL of epidermal growth factor (EGF), a potent activator of cell migration, was added. Cells were incubated for 24 hours with Δ9-tetrahydrocannabinol or cannabidiol for their ability to change bladder cancer cell migration patterns. For quantification, membranes were rinsed with cold PBS and fixed in 100% ice cold methanol for 15 min at room temperature. Fixed membranes were then stained with 0.5% crystal violet stain for 5 min to allow for visualization of the cells. Non-migrated cells were then gently removed from the upper side of the membrane with a cotton bud. Membranes were rinsed in dH_2_O until the water ran clear, allowed to dry, and then mounted on a slide. At least 3 areas of the membrane were viewed under the 10x objective of an Olympus IX81 and the number of cells for each field of view were counted. Net migration was determined by comparing the number of cells that migrated with the chemoattractant to the number of cells that migrated under control conditions (methanol).

### Invasion Assay Protocol

Growth Factor reduced 8.0 micron Matrigel Invasion Chambers (Corning) and Cell Culture Inserts with a 8.0 micron membrane were added to a 24 well plate. Matrigel Invasion Chambers were hydrated with 250 μl of medium containing 0.2% Fetal Bovine Serum, Penicillin-Streptomycin, for one hour at 37°C. T24 cells were then seeded in DMEM without FBS at a concentration of 100,000 cells/mL. 700 μl of DMEM containing 10% FBS was added to the lower chamber of invasion chambers and cell culture inserts. 250 μl of DMEM containing 0.2% FBS and 2.5 μM of Δ9-tetrahydrocannabinol, cannabidiol or vehicle control was added to the cell culture inserts and then 250 μl of the cell suspension was added to each cell culture insert and Matrigel invasion chamber. After 24 hours, media and cells that did not migrate were removed from the inside of the insert. Wells were placed in Methanol for 10 minutes and then transferred into a 3.5 g/L Crystal Violet in 2% ethanol solution for 10 minutes. Wells were then rinsed with H_2_0 and left to dry overnight. Cells that migrated or invaded through the membranes were counted using an Olympus CKX41 light microscope. Percent invasion was calculated by dividing the number of cells invaded in each condition by the number of cells migrated in the control.

### Assessment of synergism, additivity or antagonism

Synergies between Δ9-tetrahydrocannabinol or cannabidiol and gemcitabine, cisplatin or a combination of gemcitabine/cisplatin were studied using the checkerboard assay in T24 and TCCSUP cells. Synergy was also assessed between Δ9-tetrahydrocannabinol or cannabidiol and cannabivarin or cannabichromene. Briefly, the synergy assay was performed with 3,000 cells in 96-well plates with a final volume of 100 μL per well. Cannabinoids concentrations ranged from 0 to 10μM and gemcitabine and cisplatin concentrations between 0 and 100 μM. Fluorescence was quantified as described before using AlamarBlue^**®**^ after 48 h treatment. The analysis was performed using SynergyFinder 2.0 (28), where the Bliss independence drug interaction model was used. Drug combination responses were also plotted as 3D synergy maps to assess the potential synergy, antagonism or additive behaviours of the drug combinations. These maps provide visual representations of synergy and identified the concentrations at which the drug combinations had maximum effect on cell viability. The summary synergy represents the average excess response due to drug interactions. A synergy score of <−10 was considered as antagonistic, a range from −10 to +10 as additive and >+10 as synergistic (28,29).

### Statistical Analysis

Statistical analysis was completed using GraphPad Prism software. All error bars are representative of mean ± SEM. Unpaired student’s t-tests were performed for analysis of two independent groups. One-way ANOVA with Tukey’s post-hoc test was used to assess multi-group comparisons. *p* values are reported as follows: * *p* < 0.05, ** *p* < 0.01, *** *p* < 0.001.

## Results

### Effect of individual drugs on cell viability

Several cannabinoids were tested for the effect on the cell viability of bladder cancer cells. Cannabivarin and cannabichromene displayed some of the best effects of the cannabinoids tested, reducing cell viability of T24 cells with an IC_50_ of 5 μM and 6 μM, respectively (Fig. 1A). We also evaluated the effects of common chemotherapeutic agents; cisplatin was cytotoxic to T24 cells and TCCSUP transitional cell carcinoma cells with IC_50_ values of 10.75 μM and 6.75 μM, respectively, after 48 hours (Fig. 1B). Gemcitabine also displayed cytotoxic activity against the T24 cells with an IC_50_ of 102 nM while the response against the TCCSUP cells was not as strong with an IC_50_ of approximately 1 μM (Fig. 1C). Δ9-tetrahydrocannabinol yielded IC_50_ values of 8.5 μM in T24 cells and 13.5 μM in TCCSUP cells (Fig. 1D), while cannabidiol showed an IC_50_ of 5.5 μM in T24 cells and 10.5 μM in TCCSUP cells (Fig. 1E).

**FIG. 1.**
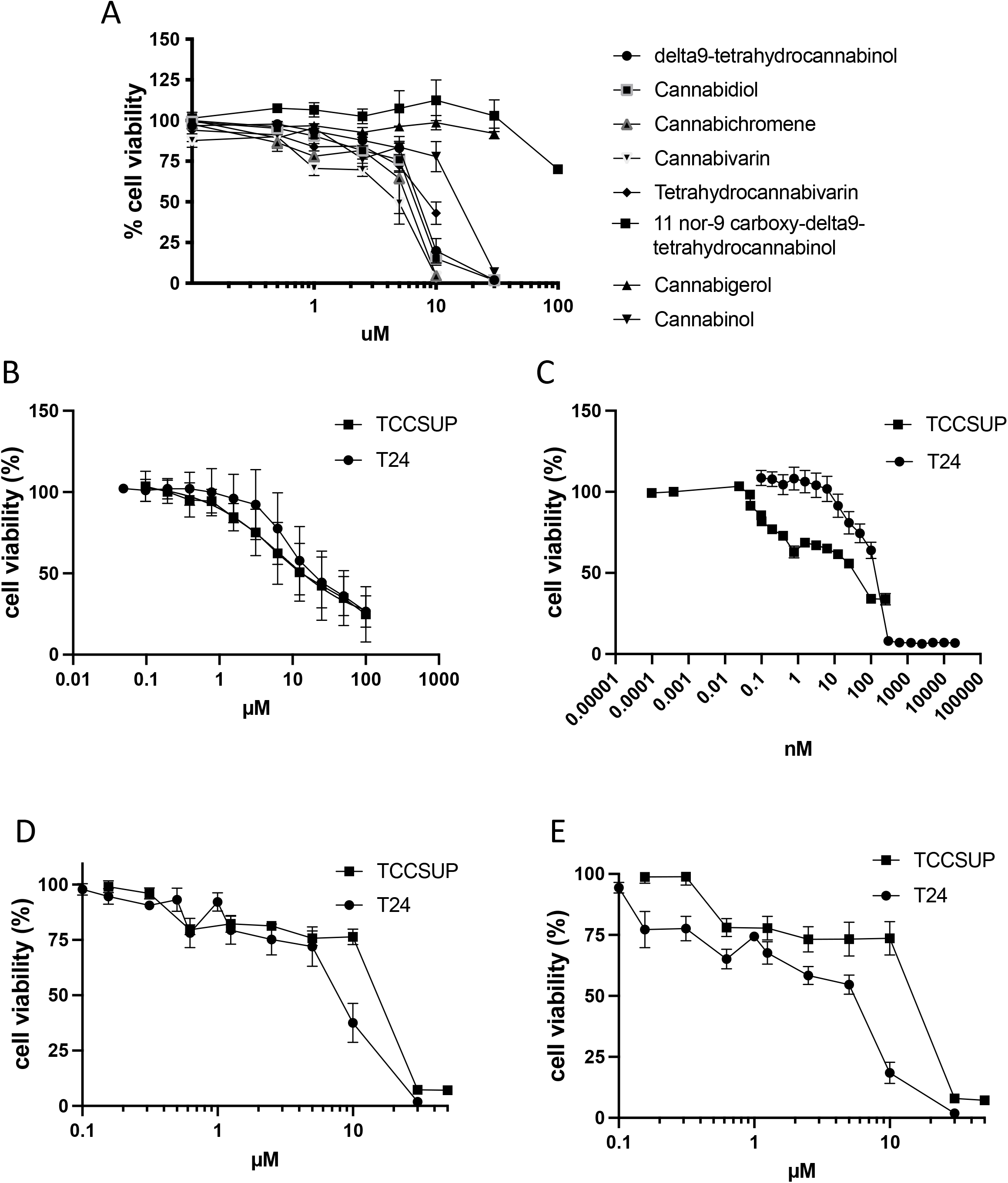
Effects of individual drugs on cell viability. Cell viability was assessed after 48 h treatment with various drugs. A. Cannabinoids; B. Cisplatin; C. Gemcitabine; D. Δ9-tetrahydrocannabinol; E. Cannabidiol. Results are means ± SEM of at least 3 independent experiments.

### Effects of cannabinoids on apoptosis

It was recently demonstrated in bladder cancer cell lines that cannabidiol and a mixture of Δ9-tetrahydrocannabinol and cannabichromene could induce apoptosis (24). In the group’s study, no results were shown regarding the effects of Δ9-tetrahydrocannabinol alone. Our results confirm the ability of cannabidiol to induce apoptosis (Fig. 2A). Following a 24 h treatment of cells with a concentration of cannabinoid at which we did not detect changes in cell viability (2.5 μM), cannabidiol induced annexin V labelling of 38.5% ± 5.5 in T24 cells and 39.3% ± 7.1 of T24 cells were labelled with annexin V following treatment with 2.5 μM of Δ9-tetrahydrocannabinol. Cannabichromene and cannabivarin at 2.5 μM also induced apoptosis with 40.6 % ± 1.6 and 41.4% ± 2.2 % of cells labelled with annexin V, respectively. Propidium iodide labelled cells following cannabinoid treatment did not differ significantly compared to the vehicle control. We investigated the potential involvement of caspase 3 in the induction of apoptosis by the two main cannabinoids Δ9-tetrahydrocannabinol and cannabidiol and observed an increase in immunoblotting of cleaved caspase 3 following ligand treatment for 24 h at a concentration of 2.5 μM (Fig. 2B).

**FIG. 2.**
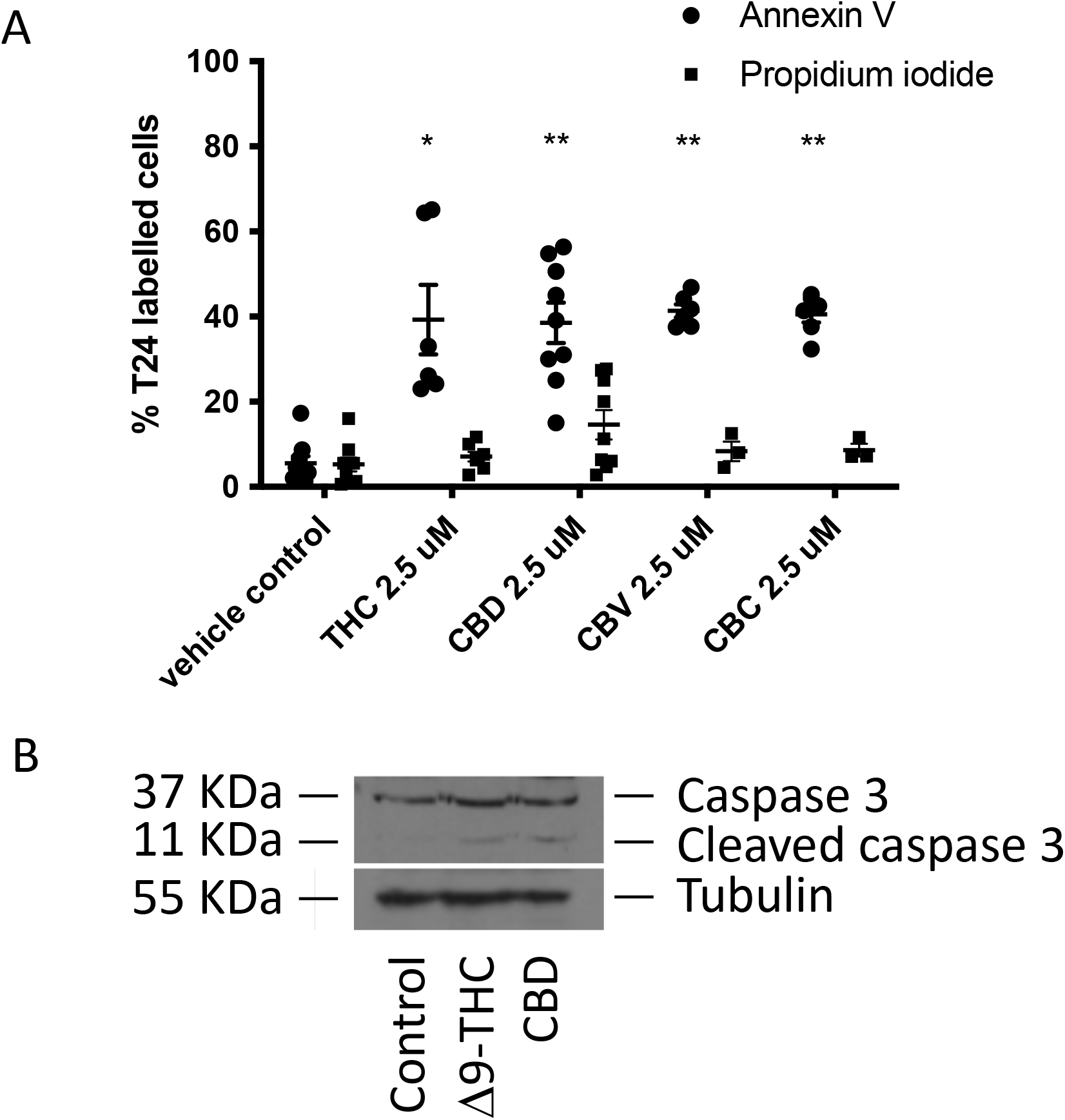
Effects of cannabinoids on apoptosis. Cells were treated for 24 h with either the methanol vehicle, Δ9-tetrahydrocannabinol, cannabidiol, cannabivarin, or cannabichromene. A. Histogram showing the % of annexin V– labeled cells and % cells stained for propidium iodide. Cells were counted from three random fields of view on a fluorescence microscope. \**p*<0.05, \*\**p* < 0.01, n = 3. B. Western blotting analysis was performed using an anti-caspase 3 antibody, and β-tubulin was included as a loading control. Figure is a representative blot of n = 3 experiments.

### Effects of cannabinoids on invasion

In addition to their anticancer effects, we evaluated the potential of Δ9-tetrahydrocannabinol and cannabidiol at reducing invasion of the high-grade and invasive T24 cells. An invasion assay was conducted where T24 cells were seeded into Matrigel invasion chambers and treated with the cannabinoids Δ9-tetrahydrocannabinol or cannabidiol for 24h. We compared the results of the Matrigel invasion chambers with chambers with migration chambers to compare migration and invasion. Our results indicate that T24 cells can invade the Matrigel (Figure 3). In our control conditions, 43.1% of cells could invade the Matrigel. Following treatment of T24 cells with 2.5 μM of Δ9-tetrahydrocannabinol for 24 hours, only 16.3% of cells could invade the Matrigel. Similarly, treatment of T24 cells with cannabidiol resulted in a reduced fraction of cells (20.2%) able to invade and cross the Matrigel.

**FIG. 3.**
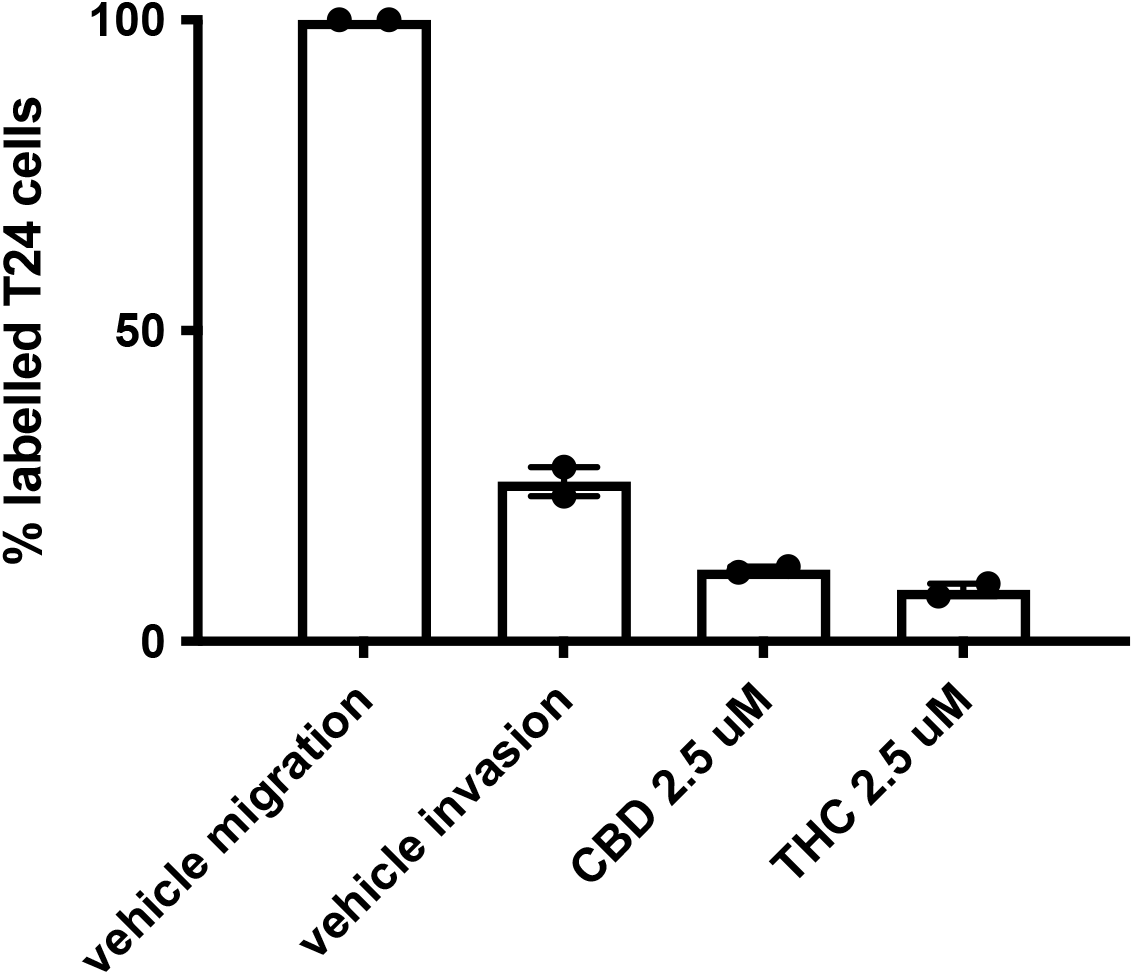
Effects of cannabinoids on invasion. Histogram summarizing Matrigel invasion assays using T24 cells in presence of either the vehicle control, Δ9-tetrahydrocannabinol or cannabidiol in comparison to vehicle treated migration. Results represent the means ± SEM of 5 experiments. \*\*\*\**p* <0.0001.

### Assessment of synergy between Δ9-tetrahydrocannabinol or cannabidiol and chemotherapeutic agents

It has been suggested that the use of combined gemcitabine-cisplatin treatment displays similar anti-cancer effects to MVAC, with fewer side effects. Therefore, we decided to test the effects of cannabinoid co-treatment with gemcitabine, cisplatin or the combination of GC on cell viability. The IC_50_ for gemcitabine and cisplatin vary from one study to another, but our values are similar to what others have previously found (30). Because of the variation in IC_50_s, we compared the ratio between gemcitabine and cisplatin among various studies and identified a range of ratios between 1:100 to 1:150 (31,32). We used a ratio of gemcitabine:cisplatin of 1:125, which is slightly above the IC_50_s we observed (1:105), but in the middle of the range of drug ratios previously published. Figure 4 shows the 3D synergy maps (28) of the combinations tested. Our results indicate that depending on the concentration of the agents used, a variety of effects can occur, from antagonism to additivity or synergy. Table 1 and Table 2 show the top and bottom 3 concentration combinations that generated the highest or lowest levels of interaction for cannabidiol and Δ9-tetrahydrocannabinol, respectively. A score <−10 is likely antagonistic; between −10 and +10 is likely additive; >+10 is likely synergistic. Our results indicate some synergy between CBD and cisplatin (Synergy score of 14) and some concentrations of CBD with gemcitabine. Some combinations between CBD and cisplatin or gemcitabine resulted in higher levels of antagonism (Table 1). Δ9-tetrahydrocannabinol displayed synergy with gemcitabine or with the gemcitabine:cisplatin combination with top synergy scores in the 14-17 range. Our results suggest that most of the effects of the combination of cannabinoids with gemcitabine or cisplatin would result in additive effects. Additive effects were observed with cisplatin, as well as some antagonism, depending on the concentrations studied.

**FIG. 4.**
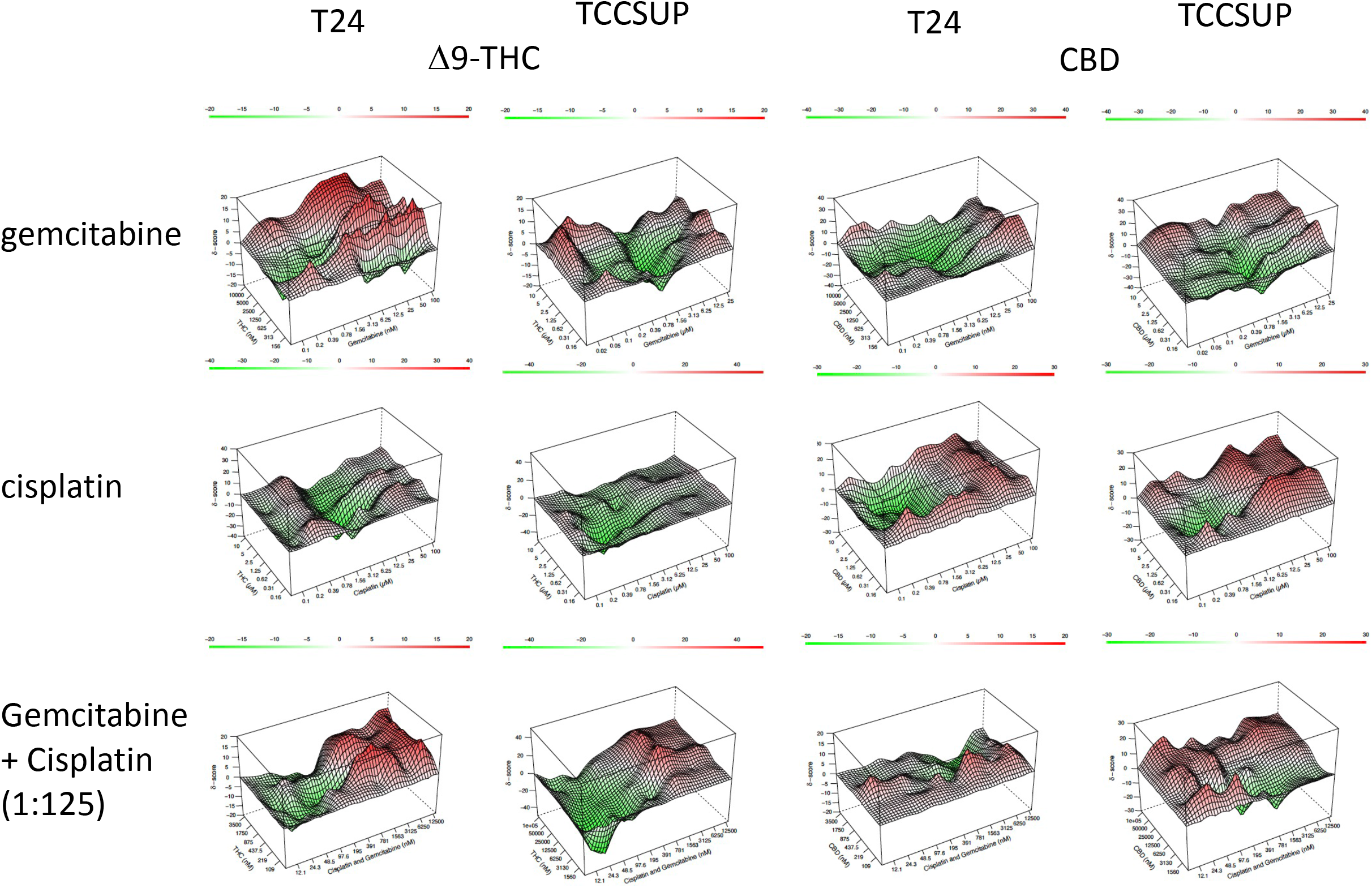
Assessment of synergy between Δ9-tetrahydrocannabinol or cannabidiol and chemotherapeutic agents. 3D synergy landscapes for the combinations of Δ9-tetrahydrocannabinol or cannabidiol with cisplatin and/or gemcitabine in T24 and TCCSUP cells.

**Table 1.**
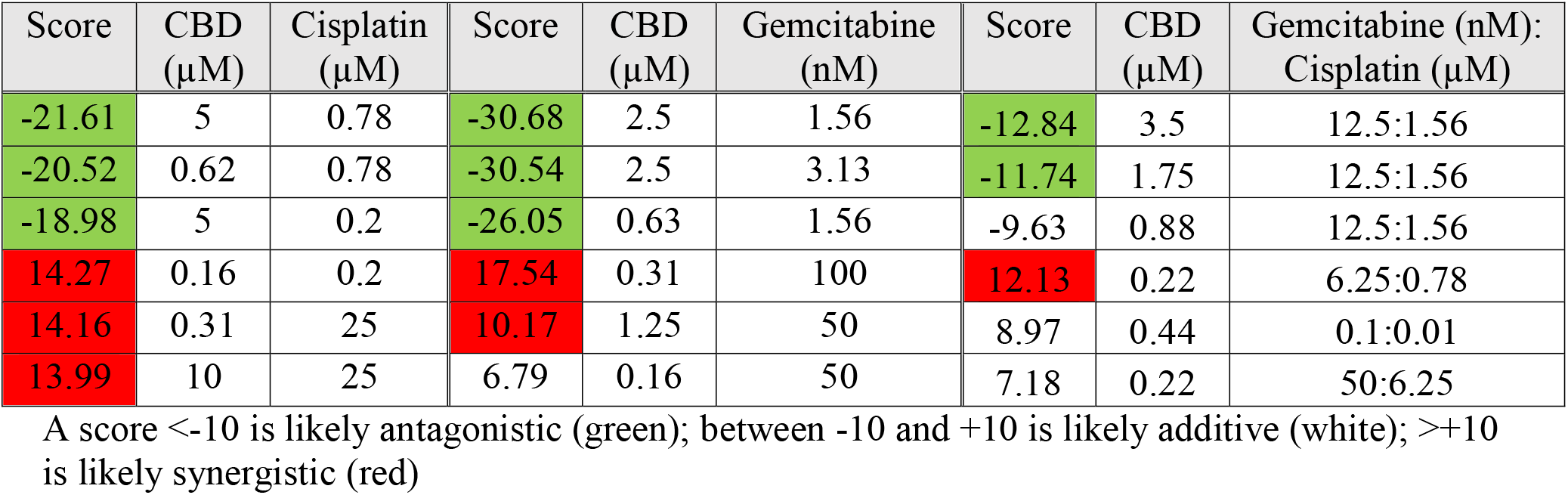
Highest and lowest levels of interaction between CBD, gemcitabine and cisplatin in T24 cells

| Score | CBD (μM) | Cisplatin (μM) | Score | CBD (μM) | Gemcitabine (nM) | Score | CBD (μM) | Gemcitabine (nM): Cisplatin (μM) |
| --- | --- | --- | --- | --- | --- | --- | --- | --- |
| -21.61 | 5 | 0.78 | -30.68 | 2.5 | 1.56 | -12.84 | 3.5 | 12.5:1.56 |
| -20.52 | 0.62 | 0.78 | -30.54 | 2.5 | 3.13 | -11.74 | 1.75 | 12.5:1.56 |
| -18.98 | 5 | 0.2 | -26.05 | 0.63 | 1.56 | -9.63 | 0.88 | 12.5:1.56 |
| 14.27 | 0.16 | 0.2 | 17.54 | 0.31 | 100 | 12.13 | 0.22 | 6.25:0.78 |
| 14.16 | 0.31 | 25 | 10.17 | 1.25 | 50 | 8.97 | 0.44 | 0.1:0.01 |
| 13.99 | 10 | 25 | 6.79 | 0.16 | 50 | 7.18 | 0.22 | 50:6.25 |

A score <-10 is likely antagonistic (green); between -10 and +10 is likely additive (white); >+10 is likely synergistic (red)

**Table 2.** Highest and lowest levels of interaction between Δ9-THC, gemcitabine and cisplatin in T24 cells

| Score | THC (μM) | Cisplatin (μM) | Score | THC (μM) | Gemcitabine (nM) | Score | THC (μM) | Gemcitabine (nM): Cisplatin (μM) |
| --- | --- | --- | --- | --- | --- | --- | --- | --- |
| -34.39 | 0.62 | 1.56 | -13.52 | 0.63 | 0.78 | -13.17 | 0.44 | 0.78:0.1 |
| -26.85 | 0.62 | 3.12 | -12.03 | 0.63 | 0.1 | -10.77 | 3.5 | 1.56:0.2 |
| -19.16 | 0.62 | 6.25 | -11.69 | 0.16 | 25 | -9.8 | 0.22 | 0.19:0.02 |
| 9.72 | 5 | 0.39 | 17.08 | 10 | 12.5 | 16.44 | 0.22 | 6.25:0.78 |
| 9.4 | 0.16 | 0.2 | 15.77 | 1.25 | 12.5 | 15.63 | 0.11 | 50:6.25 |
| 9.31 | 0.31 | 25 | 15.22 | 0.31 | 12.5 | 12.1 | 0.88 | 6.25:0.78 |

A score <-10 is likely antagonistic (green); between -10 and +10 is likely additive (white); >+10 is likely synergistic (red)

### Assessment of synergy between Δ9-tetrahydrocannabinol or cannabidiol and other cannabinoids

While the synergy results of the combinations of cannabidiol or Δ9-tetrahydrocannabinol with gemcitabine or cisplatin on cell viability were relatively low and most within the additive range, the combination of cannabinoids together produced different results. We tested the effects of the combination of Δ9-tetrahydrocannabinol or cannabidiol with cannabichromene or cannabivarin. Cannabichromene and cannabivarin reduced cell viability with better or similar IC_50_s as cannabidiol in our study. First, while some antagonism was observed, the scores were generally much closer to the additive range than what was observed with the previous set of combinations tested. The largest range of synergy scores observed occurred in the various cannabinoid combinations (Table 3). High synergy scores ranging between 38 and 71 were observed for several combinations, in particular the combinations where Δ9-THC was present (Figure 5). The combination of cannabidiol and cannabivarin did not produce synergistic effects with the same magnitude as the other cannabinoid combinations tested, with scores in the low synergistic range (between 11 and 23).

**FIG. 5.**
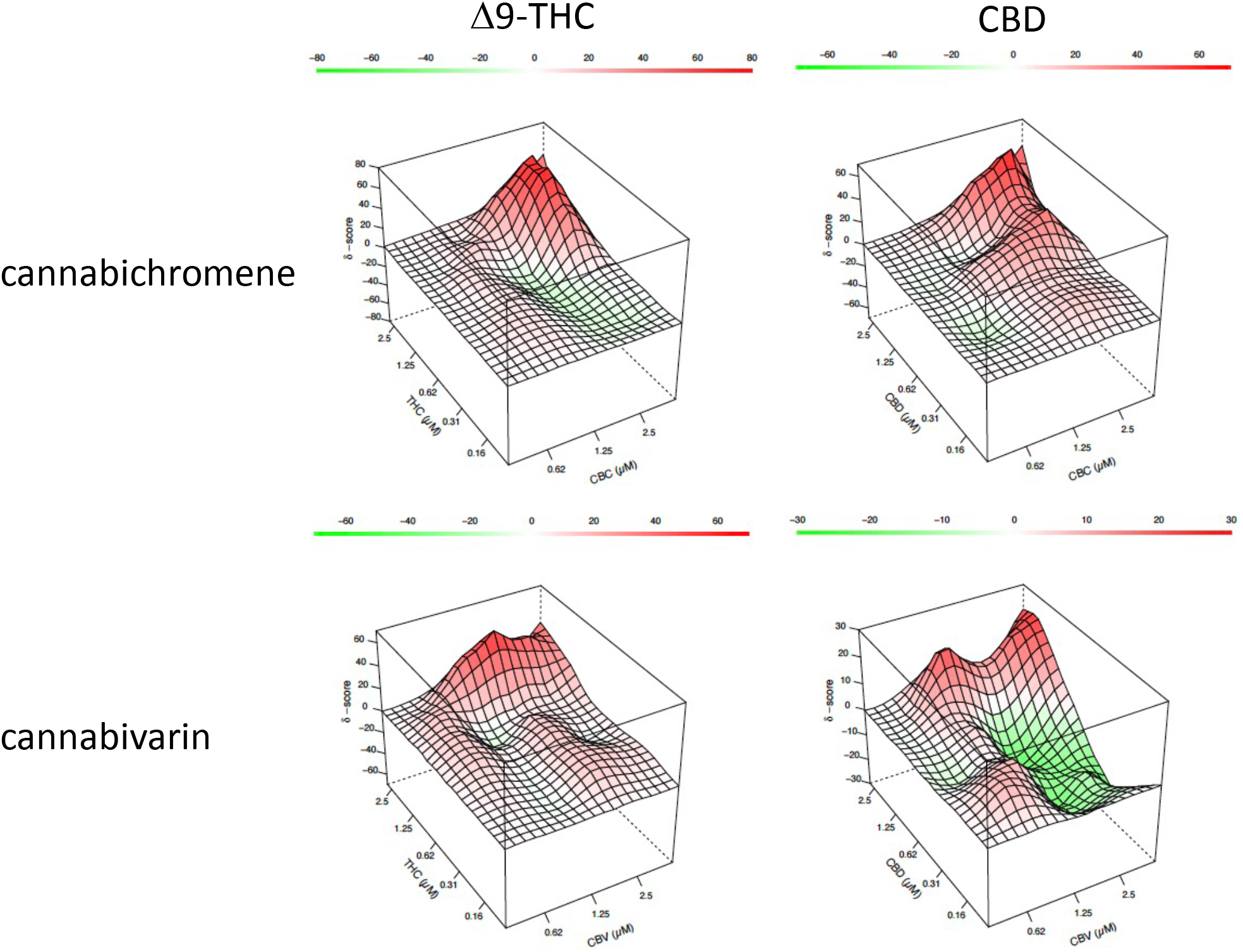
Assessment of synergy between Δ9-tetrahydrocannabinol or cannabidiol and other cannabinoids. 3D synergy landscapes for the combinations of Δ9-tetrahydrocannabinol or cannabidiol with cannabichromene or cannabivarin in T24 cells.

**Table 3.** Highest and lowest levels of interaction between Δ9-THC or CBD with CBC or CBV in T24 cells

| Score | THC (μM) | CBC (μM) | Score | THC (μM) | CBV (μM) | Score | CBD (μM) | CBC (μM) | Score | CBD (μM) | CBV (μM) |
| --- | --- | --- | --- | --- | --- | --- | --- | --- | --- | --- | --- |
| -14.41 | 0.63 | 2.5 | -8.96 | 1.25 | 1.25 | -8.22 | 10 | 10 | -20.58 | 0.31 | 10 |
| -12.56 | 0.31 | 2.5 | -5.00 | 10 | 10 | -4.55 | 1.25 | 0.63 | -18.06 | 5 | 10 |
| -10.42 | 1.25 | 2.5 | -4.74 | 0.63 | 5 | -4.55 | 2.5 | 1.25 | -18.01 | 0.31 | 2.5 |
| 42.88 | 1.25 | 5 | 40.16 | 5 | 1.25 | 38.31 | 1.25 | 5 | 11.74 | 2.5 | 10 |
| 69.34 | 5 | 5 | 45.76 | 5 | 5 | 44.64 | 5 | 2.5 | 18.65 | 5 | 5 |
| 71.43 | 2.5 | 5 | 60.03 | 5 | 2.5 | 66.35 | 5 | 5 | 23.31 | 5 | 1.25 |

A score <-10 is likely antagonistic (green); between -10 and +10 is likely additive (white); >+10 is likely synergistic (red)

## Discussion

In this study, the effects of several cannabinoids, including Δ9-tetrahydrocannabinol and cannabidiol, were tested for their potential anti-tumoral effects in bladder cancer. Our results indicate that cannabinoids can reduce cell viability of human bladder transitional cell carcinoma cell lines. We demonstrate that apoptosis is involved in the process and that caspase 3 is involved. Additionally, we show that invasion can be diminished by cannabinoids. Finally, we tested the potential synergistic effects of the combination of cannabinoids with current chemotherapeutic treatments or other cannabinoids. Our results show that some synergy occurs with gemcitabine or cisplatin, but that much higher levels of synergy are present when cannabinoids like Δ9-tetrahydrocannabinol and cannabidiol are combined with cannabichromene or cannabivarin.

Our results indicate variations in the ability of the cannabinoids tested to produce anti-tumoral effects in bladder cancer cell lines. For example, while cannabichromene, cannabivarin and cannabidiol produced effects that were generally within the same concentration range, cannabigerol, cannabinol or the metabolite 11 nor-9 carboxy-Δ9-tetrahydrocannabinol (THC-COOH) required much larger concentrations to produce an effect. We cannot dismiss that metabolites like THC-COOH could also exert anti-cancer effects, since they can accumulate at high concentrations in the urine, especially for frequent or heavy cannabis users (33). Here, we concentrated our investigation on the cannabinoids that displayed the highest levels of anti-tumour activity *in vitro*, cannabichromene, cannabivarin, cannabidiol and Δ9-tetrahydrocannabinol. The IC_50_ values observed were between 5 and 10.5 μM in T24 cells, while IC_50_ values were slightly higher in TCCSUP cells. While these levels may not be reached when cannabis is consumed, various methods including intravesical therapy, for example, could allow appropriate concentrations of the various cannabinoids to be reached to treat bladder cancer *in vivo*.

Several reports have indicated that cannabinoids may induce cell death via induction of the apoptotic cascade (34). Our results indicate that apoptosis is induced by cannabinoids and that caspase 3 is involved, as we detected cleaved caspase 3 following treatment of T24 cells with cannabinoids. These results are similar to what we and others have observed in other cancer types (23,35). Our results also indicate that not only are the apoptotic signaling pathways activated, but other signaling pathways linked to migration and invasion are also altered by cannabinoids. The invasion of high-grade and invasive T24 transitional cell carcinoma cells was reduced following treatment with cannabinoids at a concentration that did not alter cell viability. The results suggest that cannabinoids could potentially be useful to reduce migration and invasion of bladder cancer.

Multiple studies have demonstrated the ability of chemotherapeutic agents used for bladder cancer, like gemcitabine and cisplatin, to act synergistically with other compounds and produce greater anti-cancer effects (30–32). We identified that some concentrations of cannabidiol or Δ9-tetrahydrocannabinol acted synergistically with gemcitabine and/or cisplatin. These results remain to be validated *in vivo* but provide a starting point of the range of concentrations that could be required to generate effects in combination therapy involving cannabinoids.

In recent years, several studies have attempted to characterize how cannabinoids and other compounds present in the cannabis plant work together. Some have suggested an entourage effect, where the various components of the plant could work together to produce synergistic results. One study investigating the effects of pure cannabinoids versus botanical preparations has shown that their botanical preparation was more potent than pure THC at producing anti-tumor responses in both *in* vitro and *in vivo* breast cancer models (22). The compounds mediating the effects were not identified. Recently, a study demonstrated that the combination of Δ9-tetrahydrocannabinol and cannabichromene produce synergistic effects in a bladder cancer model (24). We confirmed these results, and demonstrate that other cannabinoid combinations induce synergistic effects, as observed for Δ9-tetrahydrocannabinol and cannabivarin, as well as cannabidiol and cannabichromene. Our results show the ability of different cannabinoids to produce synergistic effects when combined with other agents, including other cannabinoids.

These results remain to be validated in *in vivo* models and in human clinical trials, but overall, our results suggest that cannabinoids could potentially be useful in the treatment of bladder cancer. More investigation is needed to determine how they could be used therapeutically in the treatment of cancers, including bladder cancer, and whether as single therapy or in combination with other chemotherapeutic agents.

## Acknowledgements

The work for this study was supported by the Beatrice Hunter Cancer Research Institute Seed Funding to DJD through the Dalhousie Medical Research Foundation (DMRF) Cameron Endowment. AMT is a trainee in the Cancer Research Training Program of the Beatrice Hunter Cancer Research Institute, with funds provided by the DMRF C. MacDougall and Rosetti Funds. EGW is supported by a graduate bursary provided by the Department of Pharmacology, Dalhousie University.

## Author contributions

AMT, EGW and DJD contributed to the design of the study and performed the *in vitro* work and analysis.

## Notes

### Competing Interest Statement

The authors have declared no competing interest.

## References

1. Pons, F., Orsola, A., Morote, J., and Bellmunt, J. (2011) Variant forms of bladder cancer: basic considerations on treatment approaches. Curr Oncol Rep 13, 216–221

2. Bellmunt, J., Kim, J., Reardon, B., Perera-Bel, J., Orsola, A., Rodriguez-Vida, A., Wankowicz, S. A., Bowden, M., Barletta, J. A., Morote, J., de Torres, I., Juanpere, N., Lloreta-Trull, J., Hernandez, S., Mouw, K. W., Taplin, M. E., Cejas, P., Long, H. W., Van Allen, E. M., Getz, G., and Kwiatkowski, D. J. (2020) Genomic Predictors of Good Outcome, Recurrence, or Progression in High-Grade T1 Non-Muscle-Invasive Bladder Cancer. Cancer Res 80, 4476–4486

3. Chester, J. D., Hall, G. D., Forster, M., and Protheroe, A. S. (2004) Systemic chemotherapy for patients with bladder cancer--current controversies and future directions. Cancer Treat Rev 30, 343–358

4. Li, J., Juliar, B., Yiannoutsos, C., Ansari, R., Fox, E., Fisch, M. J., Einhorn, L. H., and Sweeney, C. J. (2005) Weekly paclitaxel and gemcitabine in advanced transitional-cell carcinoma of the urothelium: a phase II Hoosier Oncology Group study. J Clin Oncol 23, 1185–1191

5. Moore, M. J., Tannock, I. F., Ernst, D. S., Huan, S., and Murray, N. (1997) Gemcitabine: a promising new agent in the treatment of advanced urothelial cancer. J Clin Oncol 15, 3441–3445

6. Laufer, M., Ramalingam, S., Schoenberg, M. P., Haisfield-Wolf, M. E., Zuhowski, E. G., Trueheart, I. N., Eisenberger, M. A., Nativ, O., and Egorin, M. J. (2003) Intravesical gemcitabine therapy for superficial transitional cell carcinoma of the bladder: a phase I and pharmacokinetic study. J Clin Oncol 21, 697–703

7. Moore, M. J., Winquist, E. W., Murray, N., Tannock, I. F., Huan, S., Bennett, K., Walsh, W., and Seymour, L. (1999) Gemcitabine plus cisplatin, an active regimen in advanced urothelial cancer: a phase II trial of the National Cancer Institute of Canada Clinical Trials Group. J Clin Oncol 17, 2876–2881

8. von der Maase, H., Hansen, S. W., Roberts, J. T., Dogliotti, L., Oliver, T., Moore, M. J., Bodrogi, I., Albers, P., Knuth, A., Lippert, C. M., Kerbrat, P., Sanchez Rovira, P., Wersall, P., Cleall, S. P., Roychowdhury, D. F., Tomlin, I., Visseren-Grul, C. M., and Conte, P. F. (2000) Gemcitabine and cisplatin versus methotrexate, vinblastine, doxorubicin, and cisplatin in advanced or metastatic bladder cancer: results of a large, randomized, multinational, multicenter, phase III study. J Clin Oncol 18, 3068–3077

9. Bellmunt, J., von der Maase, H., Mead, G. M., Skoneczna, I., De Santis, M., Daugaard, G., Boehle, A., Chevreau, C., Paz-Ares, L., Laufman, L. R., Winquist, E., Raghavan, D., Marreaud, S., Collette, S., Sylvester, R., and de Wit, R. (2012) Randomized phase III study comparing paclitaxel/cisplatin/gemcitabine and gemcitabine/cisplatin in patients with locally advanced or metastatic urothelial cancer without prior systemic therapy: EORTC Intergroup Study 30987. J Clin Oncol 30, 1107–1113

10. Boffetta, P. (2008) Tobacco smoking and risk of bladder cancer. Scand J Urol Nephrol Suppl, 45–54

11. Freedman, N. D., Silverman, D. T., Hollenbeck, A. R., Schatzkin, A., and Abnet, C. C. (2011) Association between smoking and risk of bladder cancer among men and women. JAMA 306, 737–745

12. Thomas, A. A., Wallner, L. P., Quinn, V. P., Slezak, J., Van Den Eeden, S. K., Chien, G. W., and Jacobsen, S. J. (2015) Association between cannabis use and the risk of bladder cancer: results from the California Men’s Health Study. Urology 85, 388–392

13. Lemberger, L., Axelrod, J., and Kopin, I. J. (1971) Metabolism and disposition of delta-9-tetrahydrocannabinol in man. Pharmacol Rev 23, 371–380

14. Mehmedic, Z., Chandra, S., Slade, D., Denham, H., Foster, S., Patel, A. S., Ross, S. A., Khan, I. A., and ElSohly, M. A. (2010) Potency trends of Δ9-THC and other cannabinoids in confiscated cannabis preparations from 1993 to 2008. J Forensic Sci 55, 1209–1217

15. de Meijer, E. P., Bagatta, M., Carboni, A., Crucitti, P., Moliterni, V. M., Ranalli, P., and Mandolino, G. (2003) The inheritance of chemical phenotype in Cannabis sativa L. Genetics 163, 335–346

16. Mechoulam, R. (2005) Plant cannabinoids: a neglected pharmacological treasure trove. Br J Pharmacol 146, 913–915

17. Blázquez, C., Carracedo, A., Barrado, L., Real, P. J., Fernández-Luna, J. L., Velasco, G., Malumbres, M., and Guzmán, M. (2006) Cannabinoid receptors as novel targets for the treatment of melanoma. FASEB J 20, 2633–2635

18. Blázquez, C., Salazar, M., Carracedo, A., Lorente, M., Egia, A., González-Feria, L., Haro, A., Velasco, G., and Guzmán, M. (2008) Cannabinoids inhibit glioma cell invasion by down-regulating matrix metalloproteinase-2 expression. Cancer Res 68, 1945–1952

19. Guzmán, M., Duarte, M. J., Blázquez, C., Ravina, J., Rosa, M. C., Galve-Roperh, I., Sánchez, C., Velasco, G., and González-Feria, L. (2006) A pilot clinical study of Delta9-tetrahydrocannabinol in patients with recurrent glioblastoma multiforme. Br J Cancer 95, 197–203

20. Carracedo, A., Gironella, M., Lorente, M., Garcia, S., Guzmán, M., Velasco, G., and Iovanna, J. L. (2006) Cannabinoids induce apoptosis of pancreatic tumor cells via endoplasmic reticulum stress-related genes. Cancer Res 66, 6748–6755

21. Javid, F. A., Phillips, R. M., Afshinjavid, S., Verde, R., and Ligresti, A. (2016) Cannabinoid pharmacology in cancer research: A new hope for cancer patients? Eur J Pharmacol 775, 1–14

22. Blasco-Benito, S., Seijo-Vila, M., Caro-Villalobos, M., Tundidor, I., Andradas, C., García-Taboada, E., Wade, J., Smith, S., Guzmán, M., Pérez-Gómez, E., Gordon, M., and Sánchez, C. (2018) Appraising the “entourage effect”: Antitumor action of a pure cannabinoid versus a botanical drug preparation in preclinical models of breast cancer. Biochem Pharmacol 157, 285–293

23. Tomko, A., O’Leary, L., Trask, H., Achenbach, J. C., Hall, S. R., Goralski, K. B., Ellis, L. D., and Dupré, D. J. (2019) Antitumor Activity of Abnormal Cannabidiol and Its Analog O-1602 in Taxol-Resistant Preclinical Models of Breast Cancer. Front Pharmacol 10, 1124

24. Anis, O., Vinayaka, A. C., Shalev, N., Namdar, D., Nadarajan, S., Anil, S. M., Cohen, O., Belausov, E., Ramon, J., Mayzlish Gati, E., and Koltai, H. (2021) Cannabis-Derived Compounds Cannabichromene and Δ9-Tetrahydrocannabinol Interact and Exhibit Cytotoxic Activity against Urothelial Cell Carcinoma Correlated with Inhibition of Cell Migration and Cytoskeleton Organization. Molecules 26

25. Vallo, S., Michaelis, M., Rothweiler, F., Bartsch, G., Gust, K. M., Limbart, D. M., Rödel, F., Wezel, F., Haferkamp, A., and Cinatl, J. (2015) Drug-Resistant Urothelial Cancer Cell Lines Display Diverse Sensitivity Profiles to Potential Second-Line Therapeutics. Transl Oncol 8, 210–216

26. Martin, L. T. P., Nachtigal, M. W., Selman, T., Nguyen, E., Salsman, J., Dellaire, G., and Dupré, D. J. (2019) Bitter taste receptors are expressed in human epithelial ovarian and prostate cancers cells and noscapine stimulation impacts cell survival. Mol Cell Biochem 454, 203–214

27. Young, B., Purcell, C., Kuang, Y. Q., Charette, N., and Dupré, D. J. (2015) Superoxide Dismutase 1 Regulation of CXCR4-Mediated Signaling in Prostate Cancer Cells is Dependent on Cellular Oxidative State. Cell Physiol Biochem 37, 2071–2084

28. Ianevski, A., Giri, A. K., and Aittokallio, T. (2020) SynergyFinder 2.0: visual analytics of multi-drug combination synergies. Nucleic Acids Res 48, W488–W493

29. Ianevski, A., He, L., Aittokallio, T., and Tang, J. (2020) SynergyFinder: a web application for analyzing drug combination dose-response matrix data. Bioinformatics 36, 2645

30. Mey, V., Giovannetti, E., De Braud, F., Nannizzi, S., Curigliano, G., Verweij, F., De Cobelli, O., Pece, S., Del Tacca, M., and Danesi, R. (2006) In vitro synergistic cytotoxicity of gemcitabine and pemetrexed and pharmacogenetic evaluation of response to gemcitabine in bladder cancer patients. Br J Cancer 95, 289–297

31. Ma, Y., Yu, W. D., Trump, D. L., and Johnson, C. S. (2010) 1,25D3 enhances antitumor activity of gemcitabine and cisplatin in human bladder cancer models. Cancer 116, 3294–3303

32. Rabenstein J., F. D.-C., Hakenberg OW, Jahn D, Rutz W, Hohn A, Heuschkel M, Cuber P, Krause BJ. (2017) Monitoring cytotoxicity of gemcitabine and cisplatin in T24 bladder cancer cells by the use of F-18-FDG and F-18-FMC. Int J Clin Exp Med 10(3):4556–4564

33. Smith-Kielland, A., Skuterud, B., and Mørland, J. (1999) Urinary excretion of 11-nor-9-carboxy-delta9-tetrahydrocannabinol and cannabinoids in frequent and infrequent drug users. J Anal Toxicol 23, 323–332

34. Tomko, A. M., Whynot, E. G., Ellis, L. D., and Dupré, D. J. (2020) Anti-Cancer Potential of Cannabinoids, Terpenes, and Flavonoids Present in Cannabis. Cancers (Basel) 12

35. Rieder, S. A., Chauhan, A., Singh, U., Nagarkatti, M., and Nagarkatti, P. (2010) Cannabinoid-induced apoptosis in immune cells as a pathway to immunosuppression. Immunobiology 215, 598–605

